# Microscale sampling of the coral gastric cavity reveals a gut-like microbial community

**DOI:** 10.1101/2024.05.20.594925

**Authors:** Elena Bollati, David J. Hughes, David J. Suggett, Jean-Baptiste Raina, Michael Kühl

## Abstract

Animal guts contain numerous microbes, which are critical for nutrient assimilation and pathogen defence. While corals and other Cnidaria lack a true differentiated gut, they possess gastrovascular cavities (GVCs), semi-enclosed compartments where vital processes such as digestion, reproduction and symbiotic exchanges take place. The microbiome harboured in GVCs is therefore likely key to holobiont fitness, but remains severely understudied due to challenges of working in these small compartments. Here, we developed minimally invasive methodologies to sample the GVC of coral polyps and characterise the microbial communities harboured within. We used glass capillaries, low dead volume microneedles, or nylon microswabs to sample the gastric microbiome of individual polyps from six species of corals, then applied low-input DNA extraction to characterise the microbial communities from these microliter volume samples. Microsensor measurements of GVCs revealed anoxic or hypoxic micro-niches, which persist even under prolonged illumination with saturating irradiance. These niches harboured microbial communities enriched in putatively microaerophilic or facultatively anaerobic taxa, such as Epsilonproteobacteria. Some core taxa found in the GVC of *Lobophyllia hemprichii* from the Great Barrier Reef were also detected in conspecific colonies held in aquaria, indicating that these associations are unlikely to be transient. Our findings suggest that the coral GVC is chemically and microbiologically similar to the gut of higher Metazoa. Given the importance of gut microbiomes in mediating animal health, harnessing the coral “gut microbiome” may foster novel active interventions aimed at increasing the resilience of coral reefs to the climate crisis.

## Introduction

The gastrointestinal tract of all animals, from invertebrates to humans, hosts countless microorganisms that play an integral part in the physiology and health of their host. For example, the human gut is estimated to contain over 100 trillion bacterial cells belonging to over 1000 taxa [1], which influence all aspects of human biology, from immunity to behaviour and mental health [2, 3]. Compared to mammals, invertebrate animals such as insects often harbour less diverse gut communities [4], which nonetheless have a profound impact on their host’s fitness [5]. However, the field of gut microbiology is still in its infancy for non-model marine invertebrates. Such organisms are often very small and sometimes lack a true digestive tract, and their microbial communities are commonly characterised at the whole-organism level (i.e., bulk sampling strategy) without differentiating gastrointestinal communities from endosymbiotic or epibiotic communities [6, 7].

This bulk sampling strategy is also routinely employed for reef-building corals [8], which are sessile colonial organisms living in symbiosis with dinoflagellate microalgae (Symbiodiniaceae) that thrive from tropical to subtropical oceans. The algal symbionts provide up to 80% of the coral’s metabolic requirements via translocation of photosynthetically-fixed carbon, while the rest of the coral energy budget is met through heterotrophic feeding [9]. Prey, such as zooplankton, is digested in the gastrovascular cavity (GVC), a semi-enclosed compartment that shares many commonalities with the digestive tracts of higher Metazoa despite lacking the degree of differentiation observed in true guts [7, 10, 11]. The coral GVC is lined by endodermal tissue, and is separated from the surrounding environment by the polyp’s mouth and actinopharynx. Many central processes of holobiont physiology take place in the GVC: digestion, symbiont acquisition and expulsion, reproduction, and circulation of fluids and nutrients between inter-connected polyps [7]. Due to its morphology, the coral GVC likely presents micro-gradients not unlike those observed in bilaterian guts [7]. For example, while oxygen concentration in the external diffusive boundary layer (DBL) and in the upper GVC is primarily driven by diel light fluctuations [11, 12], a study performed on one coral species has reported a steep oxycline deeper in the GVC, leading to an anoxic zone at the bottom that can persist even under prolonged illumination [11]. Other studies have shown a pH decrease of up to one unit, as well as a decrease in the concentration of calcium ions [13, 14]. This limited microenvironmental evidence suggests that the coral GVC could be a hypoxic or even anoxic cavity, rich in carbohydrates and other metabolites from heterotrophic feeding. This would make it an ideal environment to harbour a specialised microbial community, which may play important roles in holobiont health similarly to the gut microbiome of higher metazoans.

Coral microbiomes have gained considerable attention in recent years due to their potential role in mitigating the adverse effects of ocean warming on reefs [15, 16], which causes recurrent coral bleaching events and poses the greatest threat to the survival of coral reefs [17]. To mitigate this, much research has been directed towards manipulative interventions that may increase the resilience of corals to bleaching events [18]. One of the more promising approaches involves the administration of probiotics, consortia of beneficial bacteria isolated from native coral microbiomes, which can reduce the negative effects of heat stress on the coral holobiont [19–23]. However, we still do not know how beneficial bacteria increase coral fitness [24], and more generally, what the functional role of most coral-associated bacteria is [25–28]. Microhabitat specificity is intimately linked with function [29], and communities hosted in different compartments within coral polyps (e.g., the GVC, mucus layer, tissue, skeleton) often have very different composition, functional profiles, and responsiveness to environmental change [30–34]. Bulk sampling strategies cannot identifying core bacteria that are exclusively associated with specific microhabitats (such as the algal symbiont cells) [35], an issue that hinders meaningful functional profiling. In this context, microscale sampling methods provide an invaluable tool to investigate individual microniches, including the GVC, and to unveil the role of their associated communities in holobiont health and resilience.

Technical challenges associated with sampling the coral GVC have resulted in very few attempts to characterise this specific microbiome. Using a glass microcapillary inserted through the mouth of anaesthetised polyps, Agostini et al. [11] sampled the gastric fluid from several *Galaxea fascicularis* polyps and identified a number of bacterial taxa by subcloning amplicons of 16S rDNA. Construction of a single library required pooling of approximately 0.5 mL of gastric fluid, sampled from ten polyps belonging to the same parental colony [11]. A second approach was proposed by Tang et al. [36], who collected gastric fluid from the same coral species (10-20 μL per polyp) by piercing the oral disc with a syringe and needle, subsequently plating the fluids on a rich medium (Marine Agar) and sequencing 16S rDNA from the bacterial colonies that formed. While these two approaches enabled characterisation of some GVC bacterial taxa to pioneer the study of coral GVC communities, both have limitations. Specifically, Tang et al. [36] only characterised the culturable fraction of the GVC microbiome, whilst Agostini et al. [11] avoided culturing by pooling multiple samples to obtain sufficient fluid volume. Pooling multiple samples across separate polyps not only affects the ability to analyse a large number of replicates or treatments, but also precludes the investigation of other coral species with even smaller GVCs or the characterisation of GVC heterogeneity within colonies.

Recently, a novel DNA extraction method was introduced to enable the recovery of metagenomic-quality microbial DNA from small volumes of seawater [37]. This novel method applies a physical or chemical lysis step followed by DNA recovery on paramagnetic beads to extract DNA from samples as small as 10 µL (physical lysis) or 1 µL (chemical lysis), yielding results comparable to those achieved from filtering 2 L of seawater and extracting DNA using a standard extraction kit [37]. In our present study, we therefore developed different microscale methods to sample the GVC in combination with this low-input DNA extraction protocol to characterise the microbial communities of the GVC of individual polyps for multiple coral taxa from the Great Barrier Reef (GBR). In parallel, we characterised the oxygen microenvironment experienced by these microbial communities *in hospite* using microsensors to investigate habitat specificity and potential functional profiles of the coral GVC microbiome.

## Methods

### Coral collection and aquarium maintenance

#### Great Barrier Reef (GBR) corals

Colonies of *Coelastrea aspera*, *Dipsastraea favus*, *Fungia fungites*, *Favites pentagona*, *Galaxea fascicularis* and *Lobophyllia hemprichii* (*n* = 4-6 per species, Supplementary Table S1, Supplementary Fig. 1) were collected from the reef flat of Heron Island (Great Barrier Reef, Australia) in April 2021.

#### Aquarium corals

Captive colonies of 6 genotypes of *L. hemprichii* originating from the Great Barrier Reef were obtained from the Australian ornamental trade in 2022 and maintained in aquaria at the University of Technology Sydney. Colonies were fragmented to obtain 11 sub-colonies, each with 1-3 polyps connected by tissue, yielding a total of 19 polyps (Supplementary Table S1, Supplementary Fig. 1).

Detailed information on coral sourcing and rearing conditions is provided in the Supplementary Materials (Sections 1-2).

### Micro-sensing and-sampling setup

Microsensor measurements and sampling of GVC fluid were performed in a flow chamber (Fig. 1a,b) connected to an adjustable water pump placed in a 15 L reservoir containing seawater taken from the same environment as the corals; i.e., reef water via the Heron Island Research station supply system for GBR corals, or from the UTS holding tank for aquarium corals. Flow was adjusted to ∼1 cm s^-1^, and temperature was set to 25°C with a 25W heater in the reservoir. Illumination was provided by an aquarium LED unit (Prime 16HD, Aqua Illuminations, Ames, IA, USA). A stereo microscope and/or a digital USB microscope (Dino-Lite Edge, AnMo Electronics Corporation, Taipei, Taiwan) enabled visualisation of the coral polyp mouth (Fig. 1b). Prior to performing microsensor profiles on each polyp, the bottom of the gastric cavity was identified by inserting a thin (∼75-100 µm wide) glass capillary mounted in a micromanipulator (MM33; Märzhäuser GmbH, Germany) and recording the depth at which it flexed slightly.

**Figure 1.**
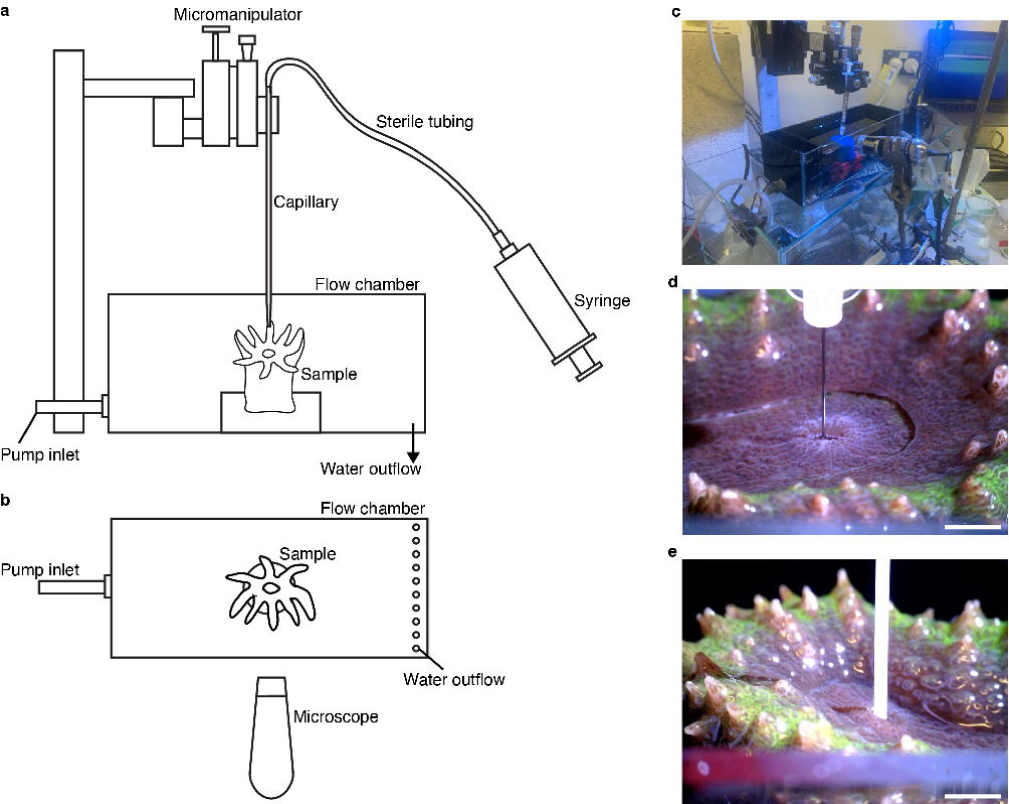
The gastric cavity sampling set up. Side (**a**) and top (**b**) view schematic illustration of the microcapillary sampling set up. (**c**) Microneedle sampling set up. (**d**, **e**) Representative images of *L. hemprichii* during sampling with a microneedle (**d**) and a microswab (**e**). Scale bars = 5 mm.

### Gastric cavity fluid sampling

#### Capillary method

GVC fluid extraction of GBR corals was performed with glass capillaries ∼75-100 µm diameter, produced by pulling glass Pasteur pipettes on a flame. The capillary was mounted on a micromanipulator (Fig. 1a) and connected to a 50 mL syringe via silicone tubing. Prior to sampling, the capillary was sterilised with 10% bleach and 80 % ethanol, then rinsed with Milli-Q water. The capillary was preloaded with Milli-Q water, which was released to equalise the pressure inside the flow chamber once the desired sampling depth was reached. After equalisation, the capillary was moved to just above the polyp mouth using the micromanipulator, then lowered into the GVC to 50% of the polyp depth before slowly collecting ∼20-50 µL of fluid over 45-60 s. The fluid was collected into a 1.8 mL cryovial (CryoPure, Sarstedt, Nürnberg, Germany) and homogenised by pipetting. A detailed sampling protocol including all sterilisation and equalisation procedures is provided in the Supplementary Materials (Section 3).

Immediately after homogenisation, a 5 µL subsample was fixed in 2% glutaraldehyde in 3× PBS (final volume 100 µL) for flow cytometry analysis, incubated for 20 minutes and then snap frozen in liquid nitrogen. The remaining fluid (typically 15-30 µL total volume depending on polyp size) was snap-frozen immediately for later DNA extraction. Three polyps per species were sampled with this method (except for *F. fungites*, a non-colonial coral, for which only a single polyp was sampled). The same sampling approach was used to collect water samples from the diffusive boundary layer (DBL) of each coral, about 30-50 µm above the oral disk surface and equidistant between the mouth and the polyp/corallite wall, and from the overlying seawater.

#### Needle method

Needle sampling of GVC fluid was performed on aquarium *L. hemprichii* polyps using a sterile low dead volume needle (34G, 9 mm long; The Invisible Needle, TSK, Vancouver, BC, Canada) connected to a 1 mL Luer lock syringe (Fig. 1c,d). Each coral was positioned so that the mouth opening was as close as possible to the water surface, while keeping the entire animal submerged, in order to minimise the distance travelled by the needle outside the cavity. The syringe was mounted on the micromanipulator (Fig. 1c) and the needle lowered vertically into the polyp mouth using manual control. Once the needle tip disappeared fully inside the mouth (Fig. 1d), the syringe plunger was pulled very slowly in order to collect ∼100 µL of gastric fluid. Fluid was collected into a sterile (UV radiation cross-linked for 1 hour) 1.5 mL centrifuge tube and immediately frozen at -80°C.

#### Swab method

Swab sampling of the GVC of each aquarium *L. hemprichii* polyp was performed immediately after needle sampling. A nylon swab of 0.8 mm diameter (TX730, Texwipe, Kernersville, NC, USA), which had been previously sterilised (UV radiation cross-linking for 1 hour), was mounted on the micromanipulator using a plastic pipette tip (P100) as an adapter. Using the micromanipulator manual controls, the swab was lowered into the flow chamber and into the polyp mouth, where it was then moved back and forth along the x and y axes for approximately five seconds to ensure good contact with the cavity surface (Fig. 1e). The swab was then withdrawn and removed from the micromanipulator. The tip was placed inside a cross-linked 1.5 mL centrifuge tube and cut with sterile scissors, before placing the tube in a -80°C freezer. Contamination of such sampling by seawater and mucus could be minimized by lowering the water level before sampling the GVC.

### Oxygen microprofiling

Microsensor profiling was performed using a Clark-type O_2_ microsensor (OX50, 50 µm tip diameter with a slender shaft; Unisense, Denmark) in both darkness, and under a saturating photon scalar irradiance (400-700 nm) of 650 µmol photons m^-2^ s^-1^. Oxygen microsensors were calibrated at experimental temperature and salinity using air-saturated aquarium seawater (100%) and fully deoxygenated seawater (0% O_2_, achieved using a Na_2_SO_3_ solution). Prior to measurement, the coral was exposed to saturating light or darkness for 20 min to allow O_2_ concentration gradients to reach steady-state [38]. The microsensor tip was then manually positioned at the polyp’s mouth using the micromanipulator. For measurements in darkness, 20 µmol photons m^-2^ s^-1^ of green light were administered briefly to help locate the polyp mouth. Depth profiles of O_2_ concentration were measured down into the gastric cavity in vertical steps of 100 µm, with 3 s waiting time before each measurement, and a 1 s measuring period. The maximum depth limit for each profile was set to 80% of the total polyp depth measured in the respective light condition to minimise the chance of microsensor damage. Three vertical profiles were recorded consecutively for each polyp under each light condition. Three polyps per species were targeted for microprofiling (the same polyps used for gastric fluid sampling). Due to logistical issues, however, only a single polyp was successfully measured for *F. fungites*, and no polyp was successfully measured for *F. pentagona* and *G. fascicularis*. For one polyp of *L. hemprichii*, a time series of oxygen concentration was also recorded in darkness while holding a microsensor at 4 mm depth (1.2 mm from the GVC bottom) for 75 minutes.

### Bacterial cell counts

Counts of bacterial cells in the fixed gastric cavity fluid were conducted using flow cytometry (CytoFLEX LX, Beckman Coulter, USA), with filtered MilliQ water as the sheath fluid and a flow rate of 25 μL min^-1^. Fixed gastric cavity fluid was stained with SYBR Green (final concentration 1:10,000) for 15 minutes in the dark. For each sample, forward scatter (FSC), side scatter (SSC), and green fluorescence (488 nm, SYBR) were recorded [39].

### DNA extraction and 16S rDNA metabarcoding

DNA extraction from fluid samples (capillary GVC, DBL and seawater samples; needle GVC and seawater samples) was performed under a UV-clean hood using a low-input protocol (100 µL or 10 µL physical lysis extraction, Supplementary Table S1) described in Bramucci et al. [37]. All tubes and reagents (except ethanol and magnetic beads) were UV-sterilized for 1 h in a UV-crosslinker (CL-1000 Ultraviolet Crosslinker, UVP). Swab GVC and seawater samples were thawed and sonicated for 5 min at 4°C, before performing the same 100 µL physical lysis extraction protocol ensuring at each step that the buffer covered the swab tip. Swabs were removed from the tubes with a P1000 pipette before adding the magnetic beads. Extractions were performed in batches of 8 or 16 samples, and an extraction blank was included in each batch. Then, 5 µL of extracted DNA sample was used as PCR template and amplified using 16S V3-V4 primers with Illumina adapters (341F: **TCGTCGGCAGCGTCAGATGTGTATAAGAGACAG**CCTAYGGGRBGCASCAG and 805R: **GTCTCGTGGGCTCGGAGATGTGTATAAGAGACAG**GGACTACNNGGGTATCTAAT; adapters in bold) in a 30 µL reaction volume containing: 0.6 µL Velocity polymerase (Meridian Bioscience, Cincinnati, OH, USA), 6 µL Velocity buffer, 1.2 µL of each 10 µM primer, 3 µL of 10 µM dNTPs, 1 µL BSA (0.1 mg mL^-1^, final concentration) and 12 µL PCR water. The amplification cycle was 98°C for 2 min, followed by 30 cycles of 98°C:30 sec, 55°C:30 sec and 72°C:30 sec, followed by a 10 min final elongation at 72°C. Amplicons were visualised on a gel before being submitted to the Australian Genome Resource Facility (Melbourne, VIC, Australia) for indexing, sequencing on Illumina MiSeq in two separate batches (run 1 = GBR corals; run 2 = UTS aquarium corals) and demultiplexing.

### Sequencing data processing

All analysis was performed in R v4.1.1. Adaptors and primers were removed from demultiplexed reads using *cutadapt* v4.4 [40], and the *dada2* pipeline (v1.22) was then applied separately to each sequencing run in order to appropriately model the run-specific error rates [41]. Run 1 reads were truncated at 250 bp (forward) and 235 bp (reverse), while run 2 reads were truncated at 270 bp (forward) and 250 bp (reverse). The maximum number of expected errors was set to 2 for both runs.

As low-input DNA extraction methods are very sensitive to contamination, a stringent decontamination pipeline was implemented as recommended by Bramucci et al. [37]. Two extraction negatives and four PCR negatives were included in sequencing run 1, and three extraction negatives were included in sequencing run 2, along with three sampling negative controls (cross-linked MilliQ water collected near the flow chamber either via needle or swab at the end of all GVC sampling). For run 1, extraction contaminants were defined as ASVs that made up more than 0.03% of processed reads in each extraction negative control. PCR contaminants were defined as ASVs that were present in any amount in each of the PCR negative controls (with the exception of one PCR negative control, which was mislabelled and discarded). For run 2, all ASVs found in the extraction negative controls were classified as contaminants since the PCR negative control could not be sequenced. In addition, ASVs that made up more than 0.03% of processed reads in the sampling negative controls were classified as contaminants. ASV tables from run 1 and 2 were merged, and all contaminant sequences identified in either batch were removed from all samples. After removal of contaminants and negative controls, taxonomy was assigned based on the Silva database v138.1 [42] using the default *dada2* settings [41]. Sequences that were identified as mitochondria, chloroplasts or eukaryotes were removed along with any samples that had zero remaining ASVs. Additional *L. hemprichii* GVC samples (*n* = 10) which had been collected during methods optimisation were also removed from the dataset at this point. Rarefaction curves (Supplementary Fig. S2) were produced and inspected using the *vegan* v2.6-4 [43] package. As rarefaction curves indicated that sufficient sequencing depth had been achieved, no rarefaction was applied to the dataset.

### Statistical analysis

Shannon’s H index was calculated to estimate alpha diversity of GBR corals using *phyloseq* v1.42 [44], while beta diversity was assessed via nonmetric multidimensional scaling (NMDS) of Bray-Curtis dissimilarity in *vegan*. For univariate data (alpha diversity, GVC depth, cell counts), homogeneity of variance was tested via Levene’s test before applying parametric (t-test, paired t-test, ANOVA or RM-ANOVA) or non-parametric (Kruskal-Wallis) statistics. Post-hoc testing was carried out via Tukey test (following ANOVA) or Dunn’s test (following Kruskal-Wallis) when all pairwise comparisons were of interest, or alternatively via adjusted pairwise t-tests when only a selection of comparisons was of interest. Groups that contained fewer than three data points (e.g. *F. fungites*) were removed before performing any statistical analysis. Count data were square-root transformed, and proportional data were arcsine square-root transformed before applying statistics. Wherever multiple tests were performed on the same dataset, P values were adjusted using the Benjamini-Hochberg correction. Alpha was set to 0.05.

To compare beta diversity between groups, singleton ASVs were removed from the dataset, then homogeneity of dispersion was tested using *betadisper* in *vegan*. PERMANOVA was used to test for significant difference between groups, wherever dispersion was deemed homogeneous, while ANOSIM was used in cases of non-homogeneous dispersion. All multivariate tests were permuted 1000 times. For GBR corals, differentially abundant taxa were identified by aggregating data to each taxonomic level and performing GLM tests on centred log ratio (clr)-transformed data in *ALDEx2* v1.26.0 [45].

Core microbiome analysis was performed in *microbiome* v1.16.0 [46]. Core taxa for each group were identified as taxa that made up more than 0.01% of the community in more than 50% of samples for that group.

### Metagenomic predictions

Metagenomes were predicted from ASVs using PICRUSt2 v2.4.1 [47], and KO identifiers from the Kyoto Encyclopedia of Genes and Genomes (KEGG) were used to identify different predicted functions within each community [48]. This analysis was only performed on the *L. hemprichii* dataset due to the higher replication and lower dispersion in GVC community composition compared to the other GBR corals.

Seven metabolic marker genes (see Discussion for in-depth rationale) were identified from the literature [49, 50]. These were two high-affinity terminal oxidases, cytochrome c oxidase cbb3-type subunit I (*ccoN*, K00404), and cytochrome bd ubiquinol oxidase subunit I (*cydA*, K00425); two low-affinity terminal oxidases, cytochrome c oxidase aa3-type subunit I (*ctaD*, K02274), and cytochrome o ubiquinol oxidase subunit I (*cyo3*, K02298); the anaerobic transcription factor CRP/FNR family transcriptional regulator (*fnr*, K01420); the nitric oxide reductase subunit B (*norB,* K04561); and the catalase gene (*CAT*, K03781). ASVs were classified based on the presence/absence of each functional gene in their predicted metagenome, and the cumulative abundance of ASVs containing each functional gene was calculated for each sample. Taxa containing either *ccoN* or *cydA* were grouped as “High affinity”, and taxa containing either *ctaD* or *cyo3* were grouped as “Low affinity”. We emphasise that these functional profiles were based exclusively on predicted metagenomes rather than metagenomic data, therefore they do not represent the true abundance of these metabolic genes.

## Results

### Great Barrier Reef corals: gastric cavity microenvironment

To characterise the GVC of GBR corals physically and chemically, we measured GVC depth and performed microsensor measurements of oxygen concentration under saturating light and in darkness. Median GVC depth measured in the dark ranged between 0.2 mm for *C. aspera* to 6.5 mm for *L. hemprichii*, with the latter significantly deeper than in most other species (Supplementary Fig. S3) (one-way ANOVA, F_3,8_=7.46, P=0.011, followed by Tukey’ HSD test, Supplementary Table S2). GVC depth remained unaltered in the light for *L. hemprichii*, *F. fungites*, and *C. aspera*, whereas cavities contracted by 0.5 mm and 1-1.4 mm for *D. favus* and *G. fascicularis*, respectively (Supplementary Fig. S3).

Microsensor measurements showed that for most coral species examined (with the exception of *F. fungites*), O_2_ concentrations in the GVC were responsive to the light/dark cycle, with hyperoxic conditions generally detected under sustained illumination and anoxic conditions developing in the dark (Fig. 2). The GVCs of *D. favus*, *C. aspera* and *L. hemprichii* exhibited an oxycline in the light, with an anoxic region detected in the lower region even under saturating irradiance in some polyps (Fig. 2a,c,d,e). In darkness, the *D. favus* and *C. aspera* GVCs were predominantly anoxic (Fig. 2a,c,e), while *L. hemprichii* exhibited a normoxic/hypoxic region in the upper 1-2 mm of the GVC (Fig. 2e,f). *F. fungites* exhibited a unique GVC oxygen profile, with normoxic conditions maintained throughout the vast majority of the cavity regardless of illumination (Fig. 2b,f). Overall, potential permanently hypoxic or anoxic habitats were identified in the lower GVC of three out of four coral species investigated (Fig. 2e). Oxygen levels measured in the lower GVC were comparable with those reported from the lumen of mammalian hindguts, as well as different regions from invertebrate guts (Fig. 2f, Supplementary Table S3). Holding a microsensor in the hypoxic region close to the bottom of the GVC of *L. hemprichii* in darkness revealed that O_2_ concentration was not constant over time (Supplementary Fig. S4). Small fluctuations between 0 and 5 µM were observed for the first 40 minutes of darkness, followed by much larger fluctuations between 0 and 100 µM over several hours (Supplementary Fig. S4).

**Figure 2.**
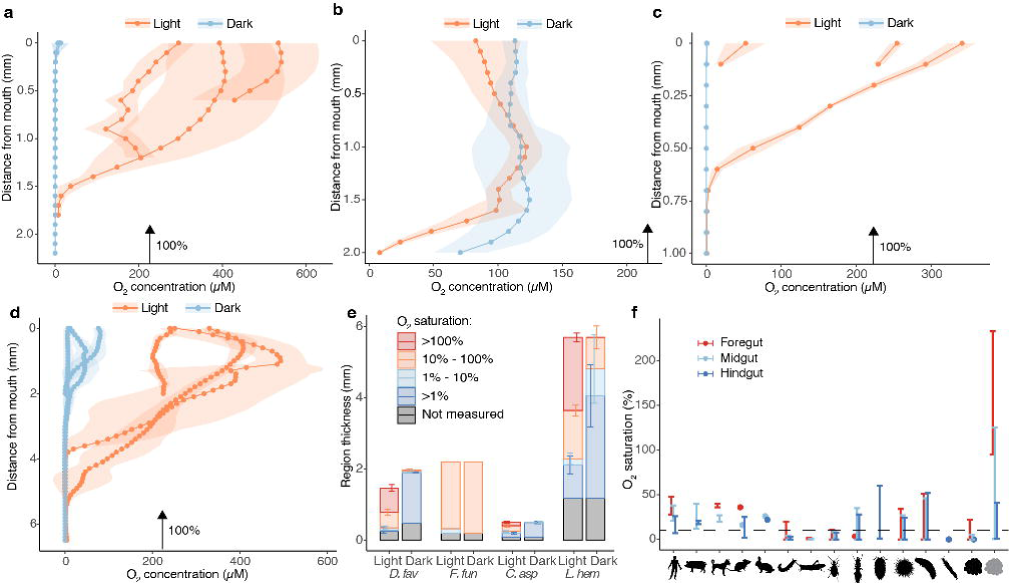
The gastric cavity oxygen microenvironment of Great Barrier Reef corals. Oxygen microsensor profiles taken inside the gastric cavity of *D. favus* (**a**), *F. fungites* (**b**), *C. aspera* (**c**), and *L. hemprichii* (**d**) collected from the Great Barrier Reef. Profiles taken under 650 µmol photons m^-2^ s^-1^ (“Light”) or in darkness (“Dark”). Arrows indicate 100% oxygen saturation under the measurement-specific temperature and salinity. Each profile corresponds to one polyp (mean ± s.d., n=3 replicate profiles per polyp). (**e**) Average thickness of GVC oxygen microniches calculated from the profiles in **a**-**d** (hyperoxic, normoxic, hypoxic and anoxic). Total bar height represents GVC depth. (**f**) Oxygen concentration ranges for different regions of the digestive tract of vertebrate and invertebrate animals (human, pig, dog, mouse, rabbit, caterpillar, grasshopper, beetle, termite, isopod, sea urchin, sea cucumber, polychaete, *L. hemprichii* in darkness, *L. hemprichii* in the light). Data for non-coral animals was calculated from the sources listed in Supplementary Table S3. The exact sections of digestive tract for each organism are listed in Supplementary Table S3 (fore, mid and hind-gut are not the technical nomenclature for all animals). For *L. hemprichii*, we considered three 2 mm thick sections of the GVC (top, middle and bottom). The partial pressure of O_2_ at sea level (21.22kPa) was considered as 100% saturation for measurements performed in air, while 100% air saturation at the measurement temperature and salinity was used for measurements performed in liquid media.

### Great Barrier Reef corals: GVC microbial community

We sampled the GVC fluid of GBR corals using the glass capillary method, and used the extracted fluid to perform bacterial cell counts and metabarcoding via 16S rDNA sequencing to characterise their gastric microbial community. Median bacterial cell counts in the GVC fluid ranged from 230,000 cells mL^-1^ (*L. hemprichii*) to 1,250,000 cells mL^-1^ (*C. aspera* and *F. fungites*), while median cell numbers in the DBL were similar across species (ranging between 420,000 in *G. fascicularis* and 614,000 cells mL^-1^ in *C. aspera*) (Fig. 3a). A significant interaction was observed between coral species and sample type (two-way ANOVA, F_6,24_=2.87, P=0.030, Supplementary Table S4); however, subsequent post-hoc pairwise t-tests did not identify specific differences between groups after adjusting for multiple testing, likely due to the small sample size (Supplementary Table S4). Median alpha diversity in the GVC (Shannon’s H index) ranged from 3.86 (*F. fungites*, single data point) to 6.81 (*C. aspera*); diversity was significantly different between groups (one-way ANOVA, F_9,28_=5.03, P<0.001), and in particular it was lower in both the *G. fascicularis* GVC and DBL compared to seawater (adjusted P<0.05 in post-hoc pairwise t-tests) (Fig. 3b, Supplementary Table S5).

**Figure 3.**
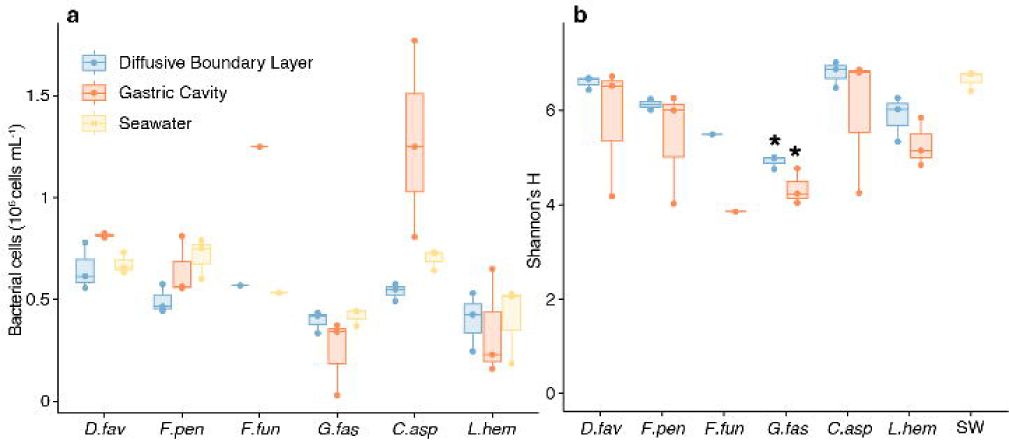
Abundance and diversity of bacteria in the gastric cavity of GBR corals. Bacterial cell counts (**a**) and alpha diversity from 16S metabarcoding (**b**) for samples collected from the GVC and DBL of *D. favus*, *F. pentagona*. *F. fungites*, *C. aspera*, and *L. hemprichii,* as well as the surrounding seawater. Spheres represent individual data point, stars show P<0.05 in Tukey’s HSD test following one-way ANOVA.

Beta diversity plots based on Bray-Curtis dissimilarity (Fig. 4a) showed that the seawater community remained similar throughout the 8-day sampling effort. Dispersion was significantly different between sampling locations (*betadisper*, 1000 permutations, F=20.04, P<0.001, Supplementary Table S5) but not between coral species (F=0.25, P=0.941) or replicate groups (F=2.27, P=0.068). Samples collected from the DBL clustered more closely together and closer to seawater, while samples collected from the GVC had greater dispersion with some replicates appearing distant not only from seawater or DBL samples, but also from other GVC samples (Fig. 4a). PERMANOVA on Bray-Curtis dissimilarity indicated that replicate groups were significantly different from each other (9999 permutations, F=2.09, R^2^=0.48 p<0.001. Supplementary Table S6).

**Figure 4.**
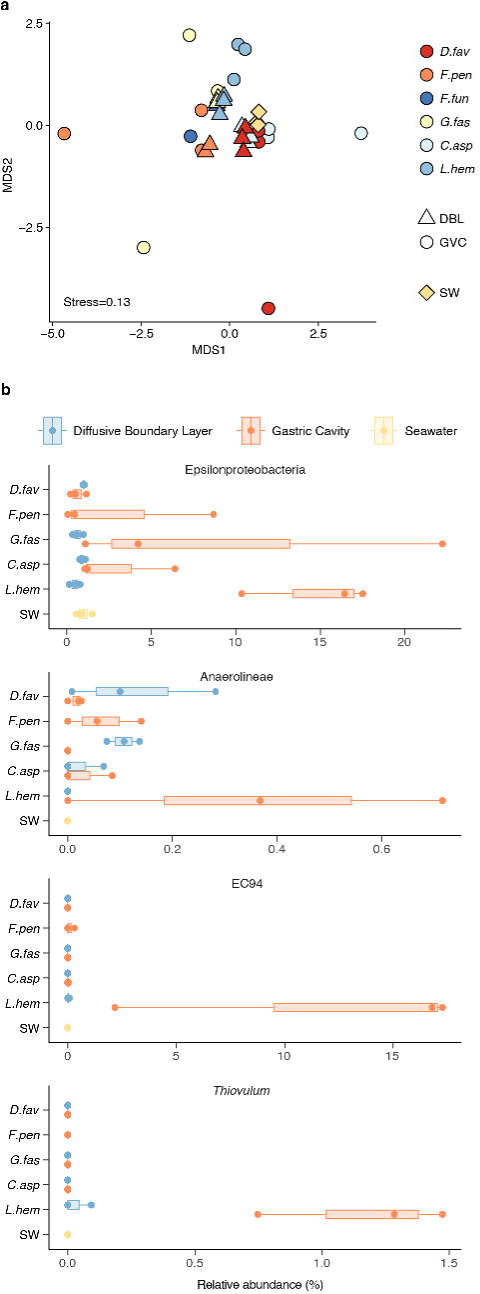
Community composition of GVC microbiomes from GBR corals. (**a**) Non-metric multidimensional scaling (NMDS) based on Bray-Curtis dissimilarity of microbial communities found in the gastric cavity (GVC) and diffusive boundary layer (DBL) of *D. favus*, *F. pentagona*, *F. fungites*, *G. fascicularis*, *C. aspera* and *L. hemprichii*, as well as seawater (SW). (**b**) Relative abundance of taxa of interest, identified as differentially abundant in coral samples by aldex Kruskal-Wallis test, in the different sample types. Spheres indicate individual data points.

Differential abundance analysis performed on taxonomically aggregated data highlighted one significantly different taxon at the phylum level (Spirochaetota, adjusted P=0.006) between coral species and sampling locations (GVC, DBL, and seawater), four at the class level (including Epsilonproteobacteria, formerly Campylobacteria, adjusted P=0.035, and Anaerolineae, adjusted P=0.048), six at the order level (including Campylobacterales, adjusted P=0.014), seven at the family level (including EC94, adjusted P=0.007), and 12 at the genus level (including *Thiovulum*, adjusted P=0.018) (Supplementary Table S7).

Differentially abundant taxa that appeared enriched in coral samples based on graphical examination are presented in Fig. 4b. Epsilonproteobacteria (formerly Campylobacteria) appeared enriched in coral GVCs, particularly in *L. hemprichii* (Fig. 4b). Anaerolineae were absent from seawater, from the *L. hemprichii* DBL and from the *G. fascicularis* GVC, but they were detected in the GVC and DBL of all other corals (Fig. 4b). The Gammaproteobacteria family EC94 was almost exclusively found in the *L. hemprichii* GVC (as well as in much smaller proportion in the *F. pentagona* GVC, Fig. 4b). The Epsilonproteobacteria genus *Thiovulum* was exclusively found in *L. hemprichii*, predominantly in the GVC as well as in very small proportion in a single sample from the DBL (Fig. 4b). Two taxa that have previously detected inside coral tissue as cell-associated microbial aggregates (CAMAs), *Endozoicomonas* and *Simkania* [51], had low abundance in the dataset. *Endozoicomonas* contributed to <1% of the community in all samples with the exception of the *F. pentagona* GVC (median = 1.8%) and DBL (median = 2.1%). *Simkania* were absent from seawater and all coral samples with the exception of a single sample of each of the following: *G. fascicularis* GVC (1.4%), *D. favus* GVC (0.76%), *F. fungites* DBL (0.84%) and *C. aspera* DBL (0.14%).

### The *L. hemprichii* GVC microbiome in aquarium and GBR corals

Next, we investigated whether core patterns in GVC microbial community composition exist across different environments (i.e., on the reef and in captivity). We sampled three additional colonies of *L. hemprichii*, then resampled all six colonies after seven days in a flow-through system with natural GBR seawater. We then compared these with GVC samples collected from aquarium colonies of the same species, which had been obtained through a commercial provider and kept long-term in an artificial seawater system.

Alpha diversity of GBR and aquarium *L. hemprichii* was significantly different between sample types (i.e., seawater; GVC and DBL of aquarium *L. hemprichii*; GVC and DBL of GBR *L. hemprichii* on the day of collection; GVC and DBL of GBR *L. hemprichii* 7 days after collection; one-way ANOVA, F_5,56_=20, P<0.001). However, post-hoc pairwise comparisons showed no significant differences between GVC communities in GBR *L. hemprichii* (whether on the day of collection or 7 days later) and in aquarium *L. hemprichii* (Fig. 5a), with the only significant differences being within samples collected from different locations (GVC vs DBL vs seawater, Supplementary Table S8). All environments tested (i.e., GVC, DBL and seawater) clearly clustered using NMDS of Bray-Curtis dissimilarity. Clustering by sample type was significant (R^2^ of 0.8; ANOSIM, 1000 permutations, P<0.001, Fig. 5b), and post-hoc pairwise comparisons confirmed that all groups were significantly different from each other (adjusted P<0.05, Supplementary Table S9). Seawater samples from the GBR formed a tight cluster, as did samples from the DBL of GBR *L. hemprichii* on the day of collection, while GVC fluid samples from both GBR and aquarium corals exhibited a wider spread (Fig. 5b). Interestingly, GVC and DBL samples from the same GBR *L. hemprichii* colonies appeared to diverge from each other 7 days after collection, and the same GVC samples clustered relatively close to those collected from aquarium *L. hemprichii* colonies (Fig. 5b). *Endozoicomonas* were absent from aquarium *L. hemprichii* samples, and *Simkania* were only detected in very low concentration (<0.1%) in two samples collected from adjacent mouths of a single individual.

**Figure 5.**
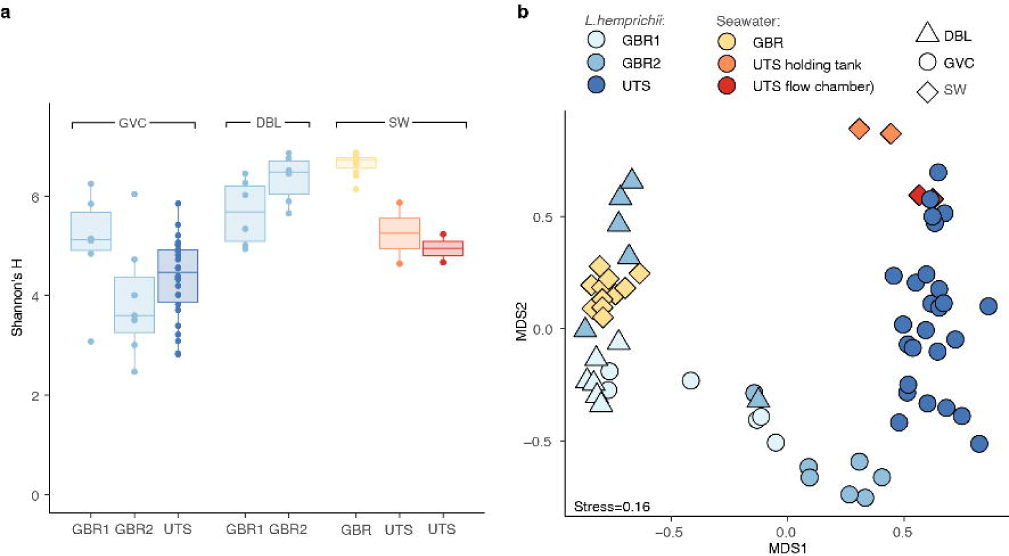
The *L. hemprichii* microbiome on the GBR and in aquarium. Alpha (**a**) and beta (**b**) diversity of microbial communities isolated from the *L. hemprichii* gastric cavity (GVC) and diffusive boundary layer (DBL) on the GBR immediately after collection (GBR1) and after 7 days in a flow-through aquarium (GBR2), from the gastric cavity of captive *L. hemprichii* (UTS), as well as from seawater, holding tanks and flow chamber. In (**a**), spheres represent individual data points.

Core microbiome analysis revealed that the DBL microbiome of *L. hemprichii* from the GBR was very variable in time (only 13.8% of the core ASVs present at the first time point were also identified as core ASVs from DBL samples at the second time point). In contrast, 64.3% of core ASVs detected in the GVC of GBR *L. hemprichii* at the first time point were also identified as core ASVs in the GVC at the second time point, and 90% of core ASVs from the second time point were also identified as core ASVs at the first time point. 68.4% of core ASVs found in the GVC of GBR *L. hemprichii* were also identified as core ASVs in the GVC of aquarium *L. hemprichii*. The 11 ASVs identified as core microbiome in both GBR and aquarium *L. hemprichii* GVC included three Epsilonproteobacteria of the order Campylobacterales, and eight Gammaproteobacteria of the family EC94. Cumulatively, these ASVs represented up to 69.0% of the bacterial relative abundance in GBR *L. hemprichii* GVC at the first sampling point (median = 18.8%), up to 83.0% when resampled (median = 50.0%), and up to 86.7% in the GVC of aquarium *L. hemprichii* (median = 14.3%) (Fig. 6a). None of these ASVs were detected in any other GBR coral or seawater sample, except for a single *L. hemprichii* DBL sample from the GBR (Fig. 6a).

**Figure 6.**
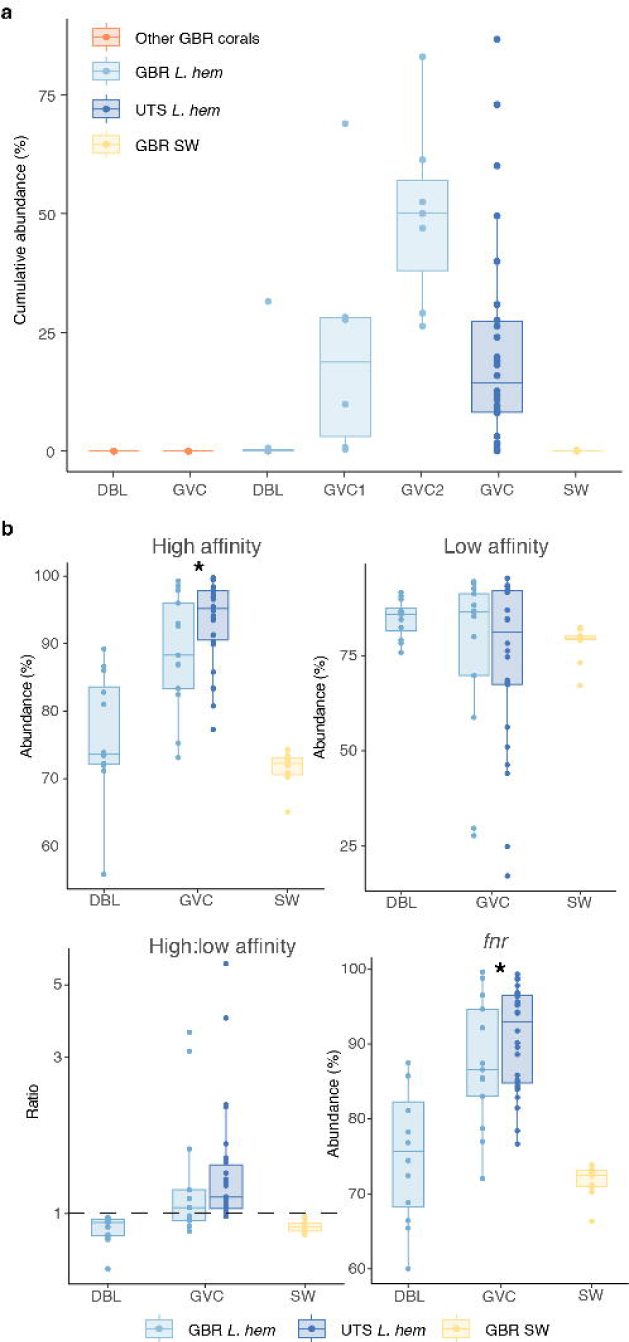
Core microbiome and predicted functional profiles in the *L. hemprichii* gastric cavity. (**a**) Cumulative abundance in different sample types of the 11 core ASVs shared by GBR and aquarium *L. hemprichii*. Sample types: gastric cavity (GVC) and diffusive boundary layer (DBL) samples collected from *L. hemprichii* (GBR *L. hem*) and other GBR corals, GVC samples from aquarium *L. hemprichii* (UTS *L. hem*), and GBR seawater samples (GBR SW). (**b**) Cumulative abundance of taxa predicted to contain the genes coding for high affinity terminal oxidases (cbb_3_ and bd type), low affinity terminal oxidases (aa_3_ and bo_3_ types), ratio between the two (y axis on log scale), and CRP/FNR family transcriptional regulator (*fnr*). Spheres represent individual datapoints. In (**b**), stars represent adjusted P<0.05 in post-hoc Dunn’s test following Kruskal-Wallis test.

Finally, we used the 16S rDNA sequencing dataset to estimate the abundance of genes that could be considered as markers of aerobic, microaerobic or (facultatively) anaerobic metabolism to investigate the potential of the *L. hemprichii* GVC to host specialised communities. Cumulative abundance of taxa predicted to contain high affinity terminal oxidases (cbb_3_ and bd types) was significantly higher in the GVC compared to the DBL and GBR seawater (Fig. 6b. Kruskal-Wallis, χ^2^=35.9, P<0.001, followed by Dunn’s posthoc test). On the other hand, no significant differences were detected in the predicted abundance of taxa containing low affinity terminal oxidases (aa_3_ and bo_3_ types. Fig. 6b. Kruskal-Wallis, χ^2^=2.27, P=0.32). The median ratio of taxa containing high:low affinity oxidases fell above 1 for GVC samples, and below 1 for DBL and seawater samples (Fig. 6b). This ratio was significantly different between groups (one-way ANOVA, F_2,59_=3.72, P=0.03), however no individual differences were highlighted by post-hoc testing (Supplementary Table S10). Taxa predicted to contain the anaerobic transcription factor *fnr* were also significantly more abundant in the GVC compared to the DBL and GBR seawater (Fig. 6b. χ^2^=32.3, P<0.001). Taxa predicted to contain the gene coding for nitric oxide reductase (*norB*) on the other hand were significantly less abundant in GVC samples compared to DBL and seawater (Supplementary Fig. S8. χ^2^=27.9, P<0.001), while those predicted to harbour the catalase gene (*CAT*) were not differentially abundant between compartments (Supplementary Fig. S8, χ^2^=0.617, P=0.735).

## Discussion

### Microscale methods to probe the gastric microbiome of reef corals

We developed and evaluated three different, yet complementary, methods to sample and characterize the gastric microbiome of corals in isolation from other compartments. Our work builds on previous attempts by Agostini et al. [11, 52], who pioneered the glass capillary method to collect gastric fluid from polyps of *G. fascicularis*. One key advancement provided by all our methods was the ability to characterise the gastric microbial community of individual polyps, eliminating the requirement to pool multiple samples in order to obtain sufficient material for molecular analysis. This was not only the case for coral species with large GVCs and large GVC fluid volumes, such as *L. hemprichii*, *F. fungites* and *G. fascicularis*, but also for species with shallower cavities and smaller fluid volumes such as *C. aspera*. Such advancement was made possible by the recent development of a low-input DNA extraction method, which enables recovery of metagenomic-quality DNA from as little as 1 μL of seawater [37]. Our approach now enables in-depth studies focusing on heterogeneity and connectivity of microbial communities at sub-colony and sub-polyp resolution, a knowledge gap previously identified by several studies of microbial diversity in coral holobionts [53–55]. In addition, our approach of sampling corals inside a flow chamber with carefully maintained environmental conditions removes the need for anaesthesia, thus enabling a closer coupling between microbial community characterisation and other physiological measurements such as O_2_ dynamics.

Our study introduced two new sampling techniques – extending beyond the glass capillary method – to target the coral gastric microbiome. Using a 34G needle to collect GVC fluid reduces the need for sterilisation of the sampling equipment, as both needles and syringes come pre-sterilised in single-use format. Such type of needle is designed to have minimal dead volume, essential when working with extremely small samples including coral gastric contents. Furthermore, the seal on the syringe plunger maintains the pressure even when the needle is lowered into or raised from the water, thus eliminating the need for complex equalisation procedures used with the glass capillary (procedures that can also lead to the loss of a small sample volume to prevent contamination). Collectively these characteristics resulted in a more streamlined, faster, and potentially more sterile sampling protocol. Sampling with a nylon microswab on the other hand aimed to target microbial taxa that might be more closely associated with the walls of the GVC, and therefore not necessarily captured when GVC fluid is collected via capillary or needle. Sampling of individual *L. hemprichii* gastric cavities with either the needle or the swab showed a relatively low overlap between bacterial taxa recovered, indicating that the two methods may indeed target different microhabitats within the cavity. However, it is unknown at this point to what extent the two methods may simply bias different microbial taxa, regardless of their location, for example through differential adherence of cells to the nylon swab, or differential release from the swab during DNA extraction [56] – this should be verified in further studies (e.g. by using appropriately constructed mock communities). Compared to the needle method, the swab sampling retrieved more unique ASVs but also more ASVs that were simultaneously detected in the surrounding seawater samples. Seawater contamination is intuitively a more substantial issue in swab samples than in needle samples since the swab is exposed while it travels through water and through the mouth before reaching the gastric cavity. To limit this issue, we recommend lowering the water level in the flow chamber as much as possible immediately prior to sampling, as even leaving the coral surface shortly exposed did not hinder insertion of the swab. We also recommend choosing carefully between the two methods depending on the specific research question, and potentially using both methods in conjunction for a more complete characterisation of the coral gastric microbiome.

### Oxygen in the coral gastric cavity: a gut-like environment?

Our characterisation of the O_2_ environment inside coral GVCs revealed some similarities between species. With the exception of *F. fungites*, all species examined presented an upper cavity environment that was generally hyperoxic in the light and normoxic or hypoxic in darkness. These characteristics are consistent with what is commonly observed in the diffusive boundary layer of corals during a diel cycle [38, 57–59]. Moving deeper into the cavity, hypoxic or anoxic regions persisted even under saturating illumination in many of the coral polyps examined. Our study thus confirms that hypoxic micro-niches, previously detected in the *G. fascicularis* GVC [11], exist in the GVC of a range of coral species. A persistently anoxic or hypoxic environment is a key feature of the digestive tract of higher metazoans including the vertebrate gut [60–63]. In fact, the combination of an anaerobic environment with a high supply of sugar is thought to be one of the factors contributing to shape gut differentiation across the tree of life [63]. Hypoxic guts support specialised microbial communities, which in many organisms contribute to the wellbeing of the host by making undigestible compounds bioavailable (termite guts represent an extreme example; [60]), by producing key metabolites (e.g. vitamins), and by defending against pathogens via antimicrobial activity [64–66]. Thus, the existence of a gut-like chemical environment in corals calls for further exploration of the microbial complement that inhabits it, and of the role these communities may play in holobiont ecophysiology.

A time series of O_2_ concentration inside the *L. hemprichii* GVC revealed that light is not the only factor shaping oxygen distribution in the GVC. Under prolonged darkness, oxygen concentrations in the GVC fluctuated from anoxic to normoxic. As no production of oxygen occurred through photosynthesis, these fluctuations were most likely due to water exchange between the hypoxic/anoxic GVC and the surrounding oxygenated seawater, possibly caused by contraction and expansion movements of the tissue that create a ventilation effect. Therefore, at least for some coral species, polyp behaviour may play a role in controlling the chemical environment of the GVC and, indirectly, the microbial community that inhabits it. The normoxic, relatively homogeneous oxygen environment of the *F. fungites* GVC could also be explained by a process of ventilation, which may be more effective in corals with larger polyps. Probing the GVC under conditions that affect tissue contraction, such as anaesthesia, stress or feeding will reveal to what extent coral polyps can regulate their GVC oxygen environment.

### The gastric microbiome of corals

Metabarcoding of microbial communities found in the DBL and GVC of GBR corals via 16S rDNA sequencing revealed that these communities are different from each other, and that they are also distinct from the surrounding seawater. While communities found in the DBL were similar to each other and similar to those found in seawater, communities sampled from coral GVCs had much wider dispersion, with some samples appearing very different not only from water samples, but also from other GVC samples. Over 50% of GVC samples from multiple species on the other hand appeared to host communities closer to the DBL and SW in composition – this was the case particularly for *D. favus*, *F. pentagona* and *C. aspera*. It is possible that potential contamination with the surrounding seawater masks the GVC community signal for certain samples only. However, it is also plausible this dispersion could result from true biological variability, whereby the GVCs of some polyps host more specialised communities while others are dominated by transient taxa found in seawater. Differences in the rate of GVC ventilation through polyp contraction, as described for *L. hemprichii*, could lead to some polyps having more extensive mixing with the surrounding environment, and therefore a microbiome that more closely resembles that of seawater or the DBL.

Intercolonial variability in microbial community composition is common across many coral taxa [67, 68], and intracolonial heterogeneity has also been previously reported when bulk sampling (i.e. combining tissue, mucus, skeleton in a single sample) [54, 69] or sampling specific compartments [53], although contrasting reports also exist [70]. Thus, GVC microbial communities found in polyps of the same species or even within the same colony could have very different composition, perhaps driven by polyp age, size, position within the colony, or recent feeding activities. Whilst this question cannot be resolved with our current dataset, the methods developed in this study are ideally suited for further investigations in this direction. Nonetheless, our data show that, at least for 30-50% of individual polyps, the GVC of all investigated GBR species hosts a microbial community that is distinct from that encountered in the surrounding seawater. The polyps with the most compositionally distinct GVC communities also exhibited lower diversity compared to the communities found in seawater. Such a notion is consistent with the observation that animal-associated microbial communities tend to have lower diversity than those found in the environment immediately surrounding them [6], and resembles what has been reported for the gut microbiome of other invertebrates, such as insects [71]. While reduced microbial diversity is an expected result in an invertebrate “gut” environment, the total number of bacterial cells retrieved from our coral GVC samples was often very similar to the cell densities recorded in seawater. This result is in contrast with a previous observation reporting two orders of magnitude more cells in the *G. fascicularis* gastric fluid compared to the surrounding seawater [11].

Metabarcoding of microbial communities found in the coral GVC highlighted a few taxa of interest. Epsilonproteobacteria (formerly Campylobacteria) were highly abundant in at least some of the GVC samples collected from all GBR coral species examined here (with the exception of *D. favus*). This group was particularly abundant in the GVC of *L. hemprichii*, including in aquarium colonies with a diverse environmental history, and some taxa of the order Campylobacterales were identified as part of the *L. hemprichii* core gastric microbiome. Epsilonproteobacteria are a class of Proteobacteria which includes many microaerophilic taxa, including known gut symbionts of other marine invertebrates [72–75], as well as mammalian gut commensals and/or pathogens [76]. Thanks to the ability of some taxa in this group to obtain energy from the oxidation of reduced compounds (chemolithotrophy) Epsilonproteobacteria dominate marine communities in sulfide-rich or hydrocarbon-rich environments, such as hydrothermal vents and sediment [76], and some taxa have become symbionts of hydrothermal vent invertebrates [77]. In corals, Epsilonproteobacteria have been previously identified as abundant taxa in tissue affected by disease or bleaching [78–81]. The presence of microaerophilic, potentially chemolithotrophic taxa in the coral gastric cavity further likens this compartment to a true animal gut, especially since some of these taxa appear to associate non-transiently with *L. hemprichii*. This discovery calls for a more in-depth investigation into the metabolism of coral gut-associated Epsilonproteobacteria to identify (i) which electron acceptors (e.g. oxygen, nitrate or sulfate) and electron donors (e.g. sulfide, thiosulfate, hydrogen) they predominantly utilise [82], and (ii) which holobiont members and physiological processes could be the source of these chemicals.

One Epsilonprotebacteria ASV found in high abundance almost exclusively in the GVC of *L. hemprichii* from the GBR was identified as *Thiovulum* sp. Members of this genus include large, highly motile sulfur-oxidising bacteria, commonly found at sulfide/oxygen interfaces where they sometimes form thick veils [83, 84]. As these cells require both oxygen and sulfide, they tend to congregate around 4% O_2_ saturation, and they are able to position themselves within the oxygen gradient via chemotaxis [84, 85]. The lower portion of the *L. hemprichii* GVC presents the ideal oxygen environment for *Thiovulum*, since this region remains hypoxic even in the light. However, a question remains regarding the potential presence and origin of sulfide in the anoxic cavity bottom, which to our knowledge has never been investigated. Sulfide production in corals has so far only been detected with microsensors under prolonged anoxic conditions, such as those that develop during exposure to organic-rich sediment [86] or infection with black band disease [87]. A similar approach could be applied to investigate the production of sulfide as well as other potential electron donors, such as hydrogen, in the GVC of healthy corals.

A second group which was more abundant in coral samples (both DBL and GVC, except for the *G. fascicularis* GVC and the *L. hemprichii* DBL) compared to seawater, and particularly abundant in the *L. hemprichii* GVC, was Anaerolineae. These are a class of Chloroflexota often isolated from microaerophilic or anoxic environments such as anaerobic digesters [88] and the mammalian gut [89], but they are also sometimes found in healthy coral tissue [81] as well as sponges [90]. This group was also reported to be enriched in seawater containing coral mucus [91]. While we cannot infer the metabolism of the specific taxa identified here simply from their taxonomic assignment, their potential involvement in fermentative pathways in the GVC is an intriguing possibility, which could have implications for digestion and resource assimilation by the holobiont.

Lastly, ASVs belonging to the family EC94 were enriched in the *L. hemprichii* GVC on the GBR, while absent from most other samples other than the GVC of *F. pentagona*. Some of these ASVs were also found in high abundance in the GVC of *L. hemprichii* from long-term aquarium culture, and were thus deemed to constitute part of the core *L. hemprichii* gastric microbiome. EC94 is a relatively uncharacterized group of marine Proteobacteria, which are predominantly associated with sponges, recently proposed for reclassification as the order Ca. Tethybacterales [92]. While members of this group are not very broadly encountered in coral samples, they appear to be dominant/core symbionts for a few coral species, including *Agaricia undata* in the Caribbean [93], *Mycedium elephantotus* in the Indo-Pacific [94], and now *L. hemprichii* on the GBR. In sponges, Ca. Tethybacterales exhibit diverse morphology and distribution, and often reside within specialized cells (bacteriocytes) [92]. Metagenome-assembled genomes (MAGs) for this group indicate they are likely aerobic or microaerophilic heterotrophs capable of utilizing a range of carbon, nitrogen and sulfur sources including dimethylsulfoniopropionate (DMSP) and glycine betaine [92], both of which are highly abundant in symbiotic corals [95].

Interestingly, we only detected low abundance of *Endozoicomonas* in the GVC of most GBR species investigated. *Endozoicomonas* are a genus of Gammaproteobacteria known to be prevalent and abundant in many coral species [96], often found as microbial aggregates (CAMAs) within the host tissue together with *Simkania* [51] – another taxon that was largely absent from our dataset. *Endozoicomonas* were also completely absent from aquarium *L. hemprichii* colonies, consistent with the common observation that *Endozoicomonas* are lost in captivity [97]. Since we did not sample the tissue directly, we cannot exclude that these corals had naturally low concentrations of these bacteria, as has been sometimes reported for corals from other locations [98]. Nonetheless, our data show that low concentrations of CAMA-forming bacteria are present also in the coral GVC, which could constitute a point of entry and exit for these microorganisms. Other potential sources of *Endozoicomonas* in the GVC include ingestion and contamination from the tissue, or resident CAMAs could exist in the GVC of corals, similarly to what observed in the gills of bivalves [99].

Alongside differential abundance and core microbiome analysis, we investigated the metabolic potential of the *L. hemprichii* and seawater microbial communities by generating predicted metagenomes and interrogating them for the presence of a set of marker genes [47, 50]. The genes coding for the terminal oxidases of respiratory chains can provide insights into the oxygen requirements of organisms [100]. Low affinity terminal oxidases include the aa_3_ and bo_3_ types, which are found in obligate aerobes and facultative anaerobes. The cbb_3_ and bd types on the other hand have a higher affinity for oxygen, thus they allow organisms to survive in low-oxygen environments (microaerophiles and some facultative anaerobes) [100]. Our analysis predicted that high affinity oxidases in the GVC of *L. hemprichii* could be (i) more abundant than low affinity ones, and (ii) more abundant than in the DBL or seawater. This suggests that the GVC may harbour a community enriched in microaerophilic and facultatively anaerobic taxa, a prediction consistent with the presence of hypoxic and anoxic zones in the lower GVC as detected by our oxygen microsensor measurements. In addition, we predicted higher abundance in the GVC for the anaerobic transcription factor gene *fnr*, which regulates the switch to anaerobic pathways in facultative anaerobes such as *E. coli* [101]. While this type of analysis is simply a prediction, if validated by metagenomic data it would provide a strong parallel with other animal gut microbiomes. High affinity terminal oxidases are the dominant (or exclusive) terminal oxidases in many vertebrate guts [100], including healthy humans [102]. Conversely, high affinity oxidases are much less abundant in environmental metagenomes, including both terrestrial and marine communities [100]. High affinity terminal oxidases are also widespread in arthropod gut microbiomes [103], including the microoxic/anoxic hindgut of termites [104].

Interestingly, the nitric oxide reductase encoding gene *norB* was predicted to be less abundant in GVC communities compared to DBL and seawater. If this prediction were to be supported with metagenomic data, it would indicate lower abundance of (facultatively) anaerobic taxa that rely on nitrate as alternative electron acceptor [49, 105]. The antioxidant enzyme catalase (*CAT*) is often used as an indicator for aerobic or oxygen tolerant species [106], as its role in detoxification of reactive oxygen species is key to survival in a high oxygen environment (however, note that some strict anaerobes also possess catalase genes [107]). We found no difference in the predicted abundance of this gene between the GVC, the DBL and seawater. We hypothesise that most taxa residing in the GVC should be able to at least tolerate oxygen, given their immediate proximity to the photosynthetic endosymbionts harboured in the coral gastrodermal tissue and given the potential ventilation occurring due to tissue contractions, which result in a highly dynamic oxygen environment. We note that predicting metagenomes from metabarcoding data can often yield misleading results due to the scarcity of annotated genomes for many bacterial taxa, as well as pervasive horizontal gene transfer occurring in microbial communities [108, 109]. However, the predicted abundance of markers *fnr*, *norB* and *CAT* has been previously shown to correlate well with metagenomic data [50]. These predictions can thus constitute a useful starting point for hypothesis generation, and can be used to guide future investigations.

## Conclusion

Multiple lines of evidence presented here highlight similarities between the coral GVC and the guts of higher vertebrates and invertebrates. The GVC contains permanently hypoxic and anoxic regions, and hosts a distinct microbial community compared to the surrounding seawater environment. The GVC community is lower in diversity and enriched in putatively anaerobic and microaerophilic taxa, including relatives of the gut microbiota of other animals. In *L. hemprichii* (the species we studied in greater detail), some of these taxa appear to form a core community which is conserved in conspecifics from different locations, and which persists after long-term aquarium culture. The microscale methods described in this article will enable further studies into the functional profiles of these communities, for example via metagenomics or metatranscriptomics, shedding light on the role played by the GVC microbiome in the physiology of the coral holobiont. We hope that these methods will pave the way towards developing “coral gut microbiology” as a new field within the broader domain of coral ecophysiological research. We anticipate that that this effort will help identify pathways and interactions within the holobiont as suitable potential targets for manipulative intervention, and eventually contribute to increasing the resilience of corals to climate change.

## Author contributions

All authors contributed to the study design. MK obtained funding. EB, DJH and JBR performed the experiments and analysed the data. EB and DJH wrote the first draft of the manuscript. All authors edited and contributed to subsequent manuscript drafts.

## Acknowledgements

We thank the staff at Heron Island Research Station for assistance during the field work, and the Great Barrier Reef Marine Parks authority for enabling our fieldwork under permit no. G18/41633.1. Caitlin Lawson for help with coral collection on Heron Island. Deepa Varkey for assistance with preliminary data analysis. Natasha Bartels, Hadley England, Kieran Chau, Nicole Dilernia and Emma Camp provided assistance with aquarium work and materials at UTS. Anna Bramucci and Trent Haydon helped with laboratory procedures at UTS.

## Data availability

All raw sequencing data has been deposited to SRA (<u>PRJNA1074944</u>). The remaining raw data is available from Dryad (doi:10.5061/dryad.p5hqbzkwj).

