## Supplementary materials for "Microscale sampling of the coral gastric cavity reveals a gut-like microbial community"

**Supplementary Methods and Results sections**

1. GBR corals collection and rearing

Fragments (~7 cm length) were collected from separate colonies of the 6 coral species using hammer and chisel, ensuring colonies were spaced a minimum of 25 m apart to avoid sampling of clones. The colonies were collected over a week and sampled on the same day as collection to capture biological and chemical properties of the GVC with minimal confounding effects arising from aquarium conditions.

Prior to sampling, colonies were maintained in indoor 30 L aquaria plumbed into a flow-through seawater system (intake located on the reef flat) at Heron Island Research Station, University of Queensland. Illumination was provided by an LED panel (Hydra 64, Aqua Illuminations, Ames, IA, USA) mounted 30 cm above the aquarium and set to provide an incident photon irradiance (400-700 nm) of ~150 µmol photons $m^{-2}$ $s^{-1}$, as measured with a small scalar irradiance sensor (US-SQS, Walz GmbH) connected to a photon irradiance (400-700 nm) meter (LI-250A, LI-COR Biosciences), using the default reef lighting profile comprising a blue-weighted spectrum and a 12h:12h light/dark cycle.

In addition to the three GBR *L. hemprichii* polyps sampled as described above, three additional polyps of *L. hemprichii* were collected from the reef to perform a heat stress experiment. These polyps were initially kept with the other specimens, sampled for GVC fluid around the same time, and finally transferred to an adjacent flow-through raceway tank (1000 L) where temperature was increased to 32°C over 7 days (at a rate of 1°C $day^{-1}$) using two custom 750 W aquarium bar heaters. Heat stressed and control GBR *L. hemprichii* polyps were then sampled again from both the DBL and the GVC with the capillary method. A third sampling point had been planned after an additional week at 32°C, however, this was abandoned after a malfunction in the research station aquarium system, which resulted in widespread coral mortality. Instead, the acquired *L. hemprichii* samples were used for comparative analysis against the aquarium *L. hemprichii* sampled at the University of Technology Sydney with the needle and swab methods.

2. Aquarium corals sourcing and rearing

Six colonies of *L. hemprichii* were obtained from the Australian ornamental trade, and identified as non-clonal due to differences in host pigment colour patterns [40]. Four colonies were obtained in June 2022 and maintained in an artificial seawater system as described in Hughes et al. [41]. Two more colonies had been obtained in January 2022 from the same supplier, and retained in a display tank under the same conditions. Fragmented colonies were divided into two groups and placed in two separate vessels in the same holding tank (700 L). Flow was provided by water exchange from the sump (via pump and gravity), heating to 25°C was provided by two 400 W heaters (Eheim, Germany), and an LED panel (Hydra 64HD, Aqua Illuminations, Ames, IA, USA) provided an incident photon irradiance (400-700 nm) of 50-80 µmol photons $m^{-2}$ $s^{-1}$, with a 12h:12h photoperiod. One group of corals ($n$ = 5 colonies) was fed daily with UV-sterilised coral pellets (Vitalis LPS, Thorne, UK), which were placed directly on the oral disk of each coral with a Pasteur pipette, while the second group ($n$ = 6 colonies) was not fed. Corals were allowed to feed for 30 minutes in the light with the pump switched off. After 30 min, any undigested pellets were removed with a Pasteur pipette. Corals were maintained in this holding tank and fed (or starved) for 15 days before GVC sampling.

3. GVC capillary sampling procedures

Glass capillaries used for sampling the GVC were produced by pulling glass Pasteur pipettes on a flame from a Bunsen burner. This resulted in elongated micropipettes with an outer tip diameter of ~75-100 µm. A fine permanent marker was used to draw two lines on the pipette corresponding to ~20 µL and ~50 µL fluid volume levels. The marked elongated pipette was then mounted on the micromanipulator stage (Fig. 1a) and sterilised by soaking it in 10% bleach for 10 min followed by thorough flushing with sterile Milli-Q water for 2 mins. A piece of silicone tubing was attached to the wide end of the capillary, while a sterile 50 mL syringe was attached to the other end of the tubing (Fig. 1a).

Prior to sampling, 50 µL of 80% ethanol was pulled into the pipette tip and held there for 1 min before flushing the pipette 10 times with sterile Milli-Q water. Then, 20 µL of Milli-Q water was drawn into the pipette tip and retained to prevent small amounts of seawater from entering while lowering the tip into the flow chamber. The micromanipulator was used to lower the pipette into the water adjacent to the coral polyp in the flow chamber. Once the desired sampling depth was reached (as previously measured), and while still outside the coral polyp, the Milli-Q water was discharged by carefully depressing the syringe so that a stream of air bubbles was observed exiting the tip. The pipette was left undisturbed until the stream of bubbles had ceased, indicating equalization of pressure with the surrounding water at that depth. Next, the pipette was very slowly raised and positioned over the centre of the polyp mouth, observing the tip very carefully to ensure that no further air bubbles were ejected, and that no seawater entered the tip. The pipette was then slowly lowered into the GVC to 50% of the polyp depth, before slowly collecting ~20-50 µL of fluid over 45-60 s. Once fluid was visible above the 20 µL marked line, the syringe was very carefully depressed so that the fluid level could be seen to be slowly dropping, while the pipette was withdrawn from the polyp and through the overlying seawater. The latter procedure was performed to create positive pressure and prevent ambient seawater from entering the pipette tip and contaminating the sample, albeit at the expense of a slightly reduced final sample volume. The sample was then collected in a cryovial, and homogenised by pipetting up and down 10 times using a P100 pipette before downstream processing.

Sampling was performed under a saturating photon irradiance (400-700 nm) of 650 µmol photons $m^{-2}s^{-1}$. as measured at the coral surface with a scalar irradiance sensor (US-SQS, Walz GmbH, Germany) connected to a photon irradiance (400-700 nm) meter (LI-250A, LI-COR Biosciences). Flow rate was measured by timing the movement of suspended particles over a given distance.

Sampling with this method had a success rate of ~25% (see next section). To minimise contamination from the surrounding seawater, polyps for which a failed sampling attempt had occurred were not sampled again. Instead, new polyps were sampled until the desired sample size was reached.

4. GVC sampling success rate

The capillary method had a success rate of ~25%, with common causes of failed sampling including: a) the inability to collect liquid into the capillary, likely indicating obstruction of the pipette tip due to particles or mesenterial filament entrainment (the most common issue encountered), b) bubbles were observed to be discharged from the Pasteur pipette tip when positioned immediately over the polyp mouth, or inside the polyp interior, which was deemed to cause mixing and risk contaminating the sample, and c) insufficient fluid volume collected (i.e. <20 µL). The success rate for needle sampling was ~60%, where the most common cause of failure was inability to draw any fluid into the needle, likely because of clogging by mucus or mesenterial filaments. Swab sampling was 100% successful; however, for one sample, no ASVs were identified following sequencing, and the sample was therefore not considered further.

5. Comparison of needle and swab sampling methods

For aquarium *L. hemprichii* colonies (sampled at UTS), we collected samples from each polyp using first the needle, then the swab method. This enabled us to directly compare the microbial community data retrieved with the two methods. Alpha diversity (measured as Shannon's H) was significantly higher for GVC samples collected with the swab compared to the needle (RM two-way ANOVA, $F_{1,22}$=14.11, P=0.004, Supplementary Fig. S5a, Supplementary Table S11), however NMDS based on Bray-Curtis dissimilarity showed no clear clustering between samples collected with the two methods (ANOSIM, P=0.462, Supplementary Fig. S5c, Supplementary Table S12). After removing ASVs that were detected in the flow chamber with either method, sampling with the swab retrieved a significantly higher proportion of unique ASVs compared to sampling with the needle (paired t-test, $t_{21}$=-2.32, adjusted P=0.041, Supplementary Fig. S6, Supplementary Table S13). However, the swab also retrieved a higher proportion of environmental ASVs, defined as ASVs that were also detected with either method from flow chamber samples ($t_{21}$=-2.60, adjusted P=0.041, Supplementary Fig. S6, Supplementary Table S13). The proportion of ASVs detected from each GVC by both methods was generally low (9.8 ± 5.3%, mean ± s.d., Supplementary Fig. S6), while the overlap between the two methods was 24.0% for the flow chamber samples, and 15.7% for the holding tank samples (note only a single sample per method was collected for flow chamber and holding tank).

### 6. Additional treatment of *L. hemprichii* colonies

As described above, some *L. hemprichii* colonies collected from the GBR were subjected to an interrupted heat stress experiment. Thus, before utilising this dataset for comparison with UTS aquarium colonies, we tested whether the treatment had a significant impact on the associated microbial communities. At the end of the interrupted heat stress treatment, only sampling location (GVC vs DBL) had a significant effect on alpha diversity (two-way ANOVA, $F_{1,9}$=18.731, P=0.0019), while the stress treatment had no significant effect ($F_{1,9}$=0.212, P=0.656) and no significant interaction was detected ($F_{1,9}$=0.004, P=0.950) (Supplementary Fig. S7a, Supplementary Table S14). NMDS of Bray-Curtis dissimilarity also showed no separation between heat stressed and control samples (PERMANOVA, $F_{1,23}$=0.78, P=0.652, Supplementary Fig. S7b, Supplementary Table S15). The GBR *L. hemprichii* samples were therefore regrouped based only on sampling day, and the dataset was used for comparison with UTS aquarium corals.

For aquarium corals, we tested for any significant effects of the feeding treatment on the microbiome before comparison with GBR corals. There was no significant difference in alpha diversity between fed and unfed corals ($F_{1,22}$=2.00, P=0.188, Figure S4b, Supplementary Table S11), and NMDS based on Bray-Curtis dissimilarity showed no clear clustering between feeding treatments (ANOSIM, P=0.099, Supplementary Fig. S5d, Supplementary Table S12). The aquarium corals were thus regrouped together, and the dataset was used for comparison with GBR corals.

**Supplementary Figures**

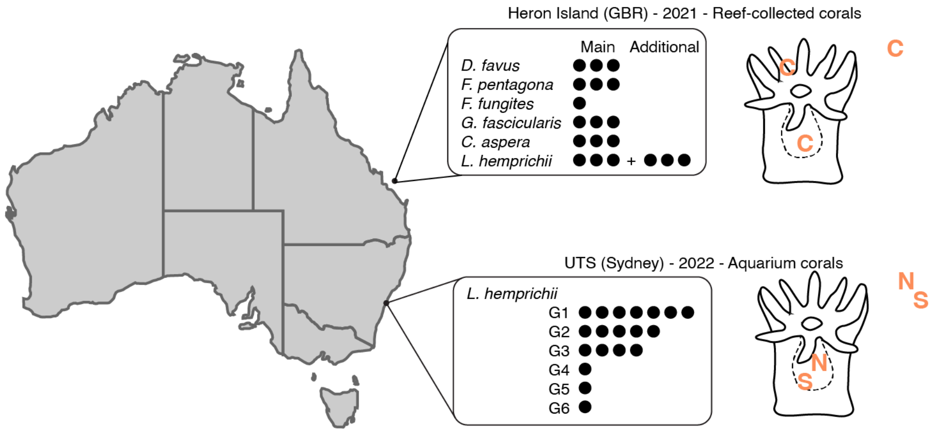


**Supplementary Figure 1. Coral origin and sampling method.** Spheres represent individual samples; G1, G2, etc. represent individual genotypes of *L. hemprichii*. Orange letters represent sampling methods C = capillary, N = needle, S = swab.

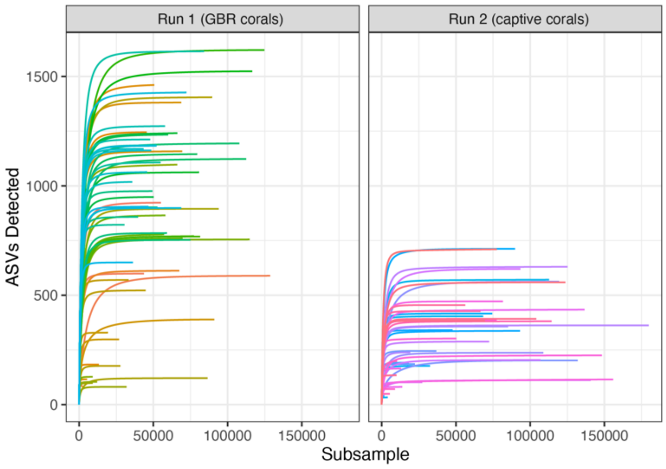


**Supplementary Figure 2. Rarefaction curves by sequencing run.**

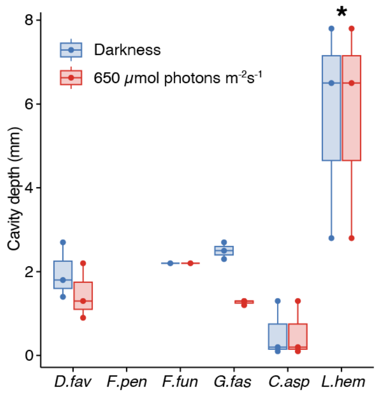


**Supplementary Figure 3. Gastric cavity depth for GBR corals.** Spheres represent individual datapoints, star represents $p<0.05$ in Tukey's test following one-way ANOVA (Supplementary Table S2).

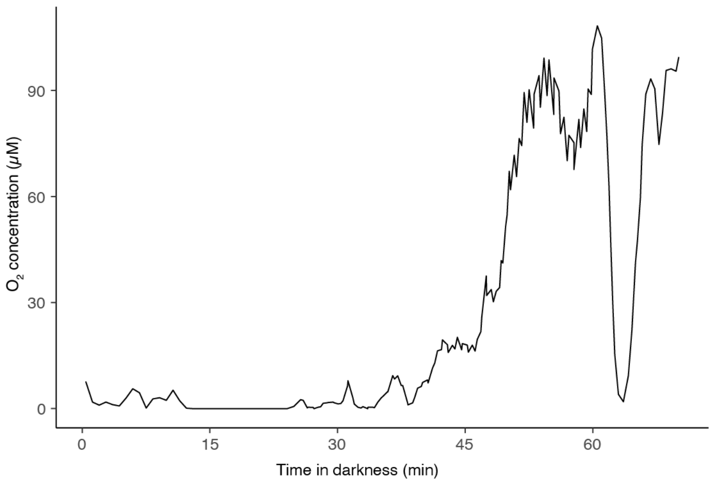


**Supplementary Figure 4. Oxygen concentration in the *L. hemprichii* GVC in darkness.** Time series obtained with a stationary microsensor placed 4mm from the polyp's mouth (estimated distance from the bottom of the cavity=1.2mm).

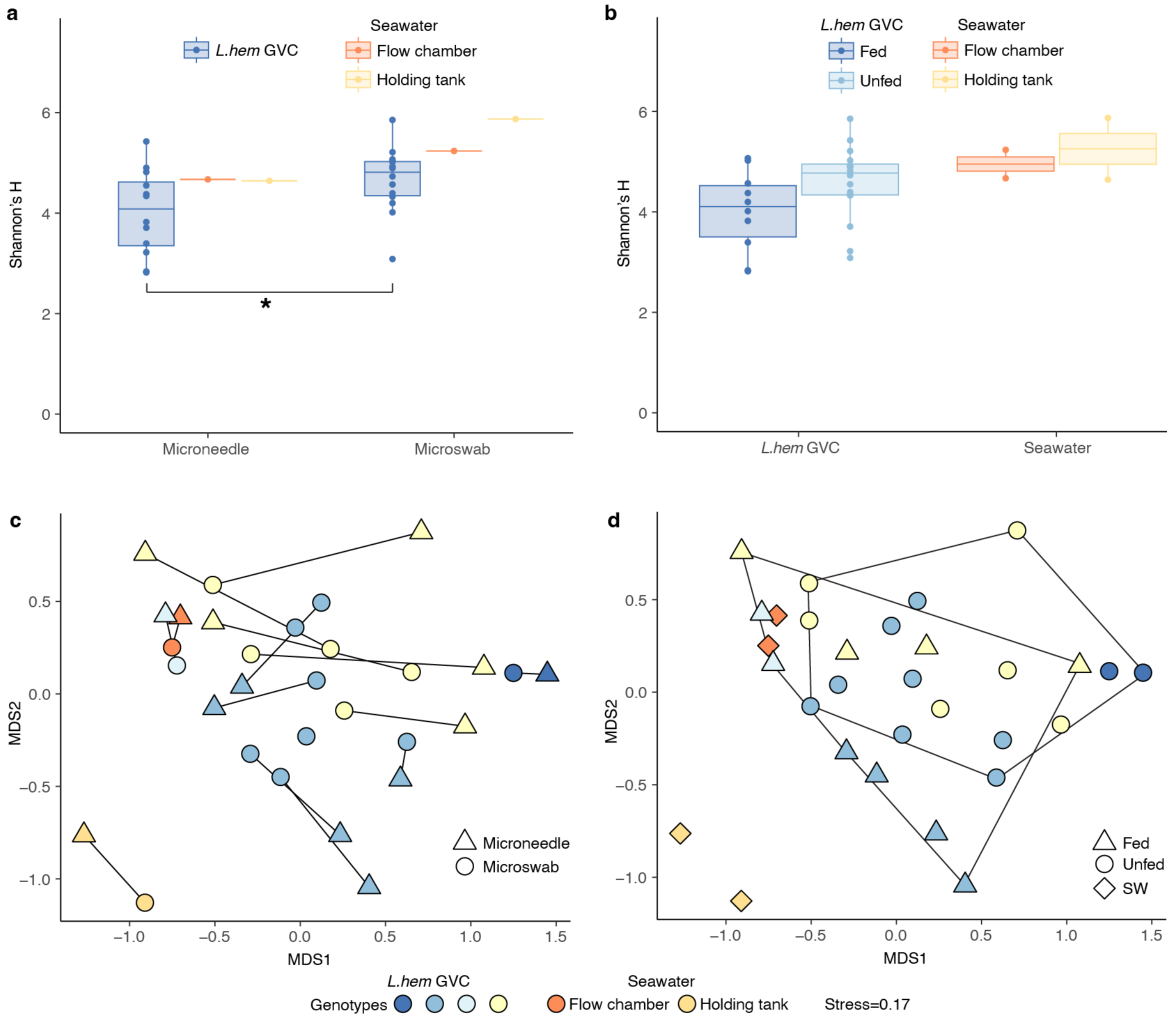


**Supplementary Figure 5. The gastric cavity microbiome of aquarium *L. hemprichii*.** Alpha (**a**,**b**) and beta (**c**,**d**) diversity of microbial communities sampled from the gastric cavity of aquarium *L. hemprichii*, grouped according to sampling method (**a**,**c**) or feeding treatment (**b**,**d**). In (**a**,**b**), spheres represent individual datapoints, star indicates adjusted P<0.05 in Tukey test following one-way ANOVA. In **c**, segments connect samples taken from the same polyp (or tank/flow chamber) with the two different methods. In **d**, polygons enclose all fed and unfed corals samples.

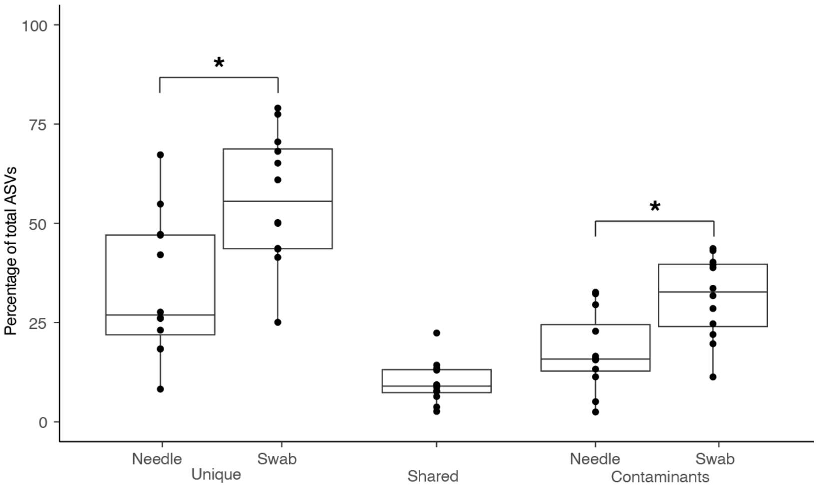


**Supplementary Figure 6. Comparison of needle and swab methods for sampling *L. hemprichii* GVC microbial communities.** Spheres represent individual samples. Stars represent adjusted P<0.05 in paired t-test.

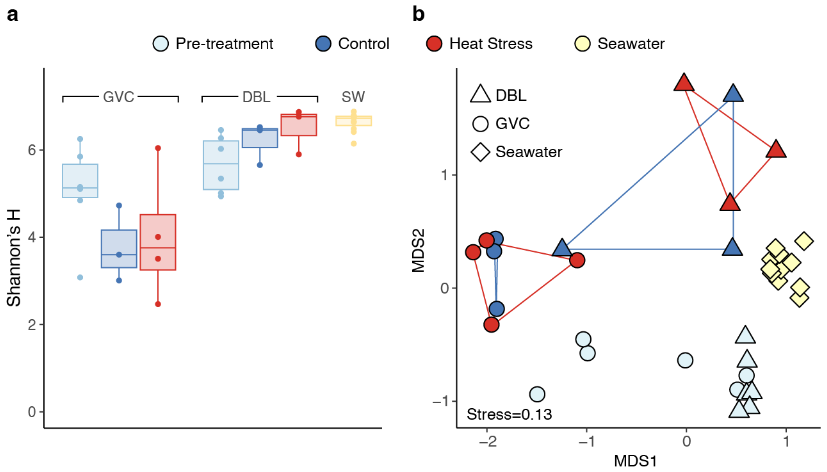


**Supplementary Figure 7. The microbiome of GBR *L. hemprichii* during heat stress.** Alpha (**a**) and beta (**b**) diversity of microbial communities sampled from the gastric cavity (GVC) and diffusive boundary layer (DBL) of *L. hemprichii* immediately after sampling from the GBR (Pre-treatment), after seven days at ambient temperature (Control), and after a 7-day temperature ramp to 32°C (Heat Stress), as well as seawater (SW). In (**b**), polygons connect samples from the GVC and DBL of control and heat stressed *L. hemprichii* after seven days.

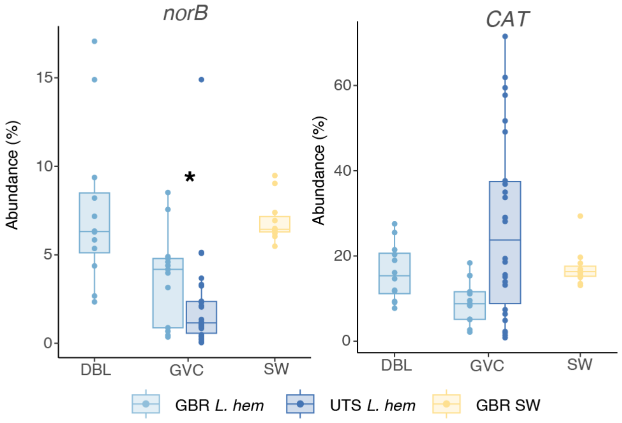


**Supplementary Figure 8. Metagenomic predictions for *L. hemprichii*.** Cumulative abundance of taxa predicted to contain the nitric oxide reductase gene (*norB*), or the catalase gene (*CAT*). Stars indicate $p<0.05$ in Dunn's test following Kruskal-Wallis.

**Supplementary Table Legends**

**Table S1.** Sample list and metadata
**Table S2.** Results of statistical analysis for GVC depth in GBR corals (Supplementary Fig. S3)
**Table S3**. Literature review of oxygen concentration in the digestive tracts of animals (Fig. 2f)
**Table S4.** Results of statistical analysis for bacterial cell counts in GBR corals (Fig. 3a)
**Table S5.** Results of statistical analysis for alpha diversity (Shannon's H) in GBR corals (Fig. 3b)
**Table S6.** Results of statistical analysis for beta diversity in GBR corals (Fig. 4a)
**Table S7.** Results of differential abundance analysis for microbial taxa associated with GBR corals, aggregated by different taxonomic levels. Differentially abundant taxa (adjusted $p<0.05$) are highlighted in blue.
**Table S8.** Results of statistical analysis for alpha diversity (Shannon's H) in *L. hemprichii* (Fig. 5a)
**Table S9.** Results of statistical analysis for beta diversity in *L. hemprichii* (Fig. 5b)
**Table S10.** Results of statistical analysis for predicted abundance of functional genes in *L. hemprichii* (Fig. 6b, Supplementary Fig. S8)
**Table S11.** Results of statistical analysis for alpha diversity (Shannon's H) in aquarium *L. hemprichii* (Supplementary Fig. S4a,b)
**Table S12.** Results of statistical analysis for beta diversity in aquarium *L. hemprichii* (Supplementary Fig. S5c,d)
**Table S13.** Results of statistical analysis for proportion of unique and contaminant ASVs sampled with the needle and swab methods in aquarium *L. hemprichii* (Supplementary Fig. S6)
**Table S14.** Results of statistical analysis for alpha diversity (Shannon's H) in GBR *L. hemprichii* at the end of the heat stress experiment (Supplementary Fig. 7a)
**Table S14.** Results of statistical analysis for beta diversity in GBR *L. hemprichii* at the end of the heat stress experiment (Supplementary Fig. 7b)
